# Probing cortical excitability under GABAergic modulation

**DOI:** 10.1101/2021.02.18.431873

**Authors:** Grégory Lepeu, Ellen Van Maren, Kristina Slabeva, Markus Fuchs, Juan Anso, Werner J. Z’Graggen, Claudio Pollo, Kaspar A. Schindler, Antoine Adamantidis, Maxime O. Baud

## Abstract

Cortical excitability, the variable response to a given cortical input, is widely studied in neuroscience, from slice experiments and in silico modeling work to human clinical settings. However, a unifying definition and a translational approach to the phenomenon are currently lacking. For example, at the onset of epileptic seizures, cortical excitability may impair resilience to perturbations (external or endogenous). In this study, we tested *in vivo* whether changes in cortical excitability quantified as evoked response to small perturbation corresponded to changes in resilience to larger perturbations. To do so, we used both cell-type circuit specific optogenetic stimulation in mice and direct intracranial stimulation in one human subject and quantified 1) evoked cortical responses to single pulses of varying intensity, and 2) evoked cortical facilitation and suppression to paired pulses at varying intervals. In the presence of a gamma-Aminobutyric acid (GABA) agonist or antagonist, we found that 1) cortical response to single pulses and 2) cortical facilitation decreased and increased, respectively. Additionally, using trains of opto-pulses in mice in the presence of a GABA agonist, we found increased resilience to the induction of seizures. With this study, we provide evidence for a tight correlation between cortical excitability and resilience, exploring a range of cortical dynamics, from physiological excitability, to pathological discharges. Our study carried out with two different stimulation methods in two species suggests that varying cortical excitability can be tracked with simple protocols involving minute short-lived perturbative stimuli.

## Introduction

The cerebral cortex is the essential information processing system of the brain, which underlies behaviors and cognitive functions in mammals. To fulfil these functions, the cortex must be excitable to respond to incoming stimuli and pass on the information, but also resilient to avoid that strong external stimuli or endogenous stochastic perturbations disrupt its dynamical equilibrium. Many cortical disorders, including epilepsy^1^, migraine^2^ or schizophrenia^3^ are believed to result from specific imbalance in this fine equilibrium. As a matter of facts, many drugs widely used to treat these disorders, such as anti-epileptic drugs, neuroleptics and benzodiazepines, act on the cortex by decreasing excitability and/or increasing resilience.

In this context, cortical excitability is defined as the variable response to incoming stimuli, whereas resilience is defined as the system’s capacity to promptly recover from perturbations^4–7^. In any mammalian brain, when perturbations overcome cortical resilience, frankly abnormal cortical dynamics arise, resulting in a self-sustained but transient seizure^4,8–10^. Defining and quantifying the boundaries between physiological and pathological cortical dynamics remains an active field of research, which has recently included the formalism of dynamical systems theory to characterize ictal transitions (i.e. transitions to seizures)^4,5,11,12^. Specifically, recent modeling and experimental work suggested that such transition occurs by the alignment of small stochastic perturbations and of a slower decrease in cortical resilience^4,11^, also called the slow permittivity variable^10,11^. However, this work mostly relied on measurements of excitability and resilience in *in-vitro* models of epilepsy, and *in-vivo* validation of these concepts and their broader applicability to the non-epileptic brain are currently lacking.

From a translational perspective, the ability to track cortical excitability over time in chronic dynamical brain disorders such as epilepsy and other brain conditions, may open the way to refined diagnostic and therapeutic approaches. In current clinical settings, invasive EEG recordings for presurgical workup of epilepsy open the possibility of delivering minute direct electrical stimulations to the cortex, to probe cortical excitability in different part of the brain, and better delineate epileptic parenchyma^13–15^. Combining this diagnostic approach with the use of anti-epileptic drugs could help better understand cortical excitability and its modulation by pharmacological agents.

In neuroscience research, cortical excitability and drug effects have long been measured as the evoked response to an external stimulus such as transcranial magnetic stimulation in humans^16–19^ or direct cortical electrical stimulation in animals^20^. These methods are, by design, not specific for circuit or cell types and may alternatively excite or inhibit the cortex depending on the specific stimulation protocol used^16^. More recently, the development of optogenetics in animal research^21^ has afforded the long-needed cell-type specific probing of brain circuits, opening the possibility to characterize cortical excitability more specifically. Optogenetics has also been used to induce seizures in healthy animals, circumventing the specific limitations of individual epilepsy models^22–24^.

At the cellular level, animal studies have now shown that, tight regulation of cortical excitability is governed by an interplay between excitatory and inhibitory neurons and that disruption of this balance can lead to epileptic seizures^25^. In particular, GABAergic inhibition plays a key role in regulating cortical excitability^26–28^ as well as seizure likelihood^29^, threshold^24^, duration^30^ and severity^31^. Further, GABAergic transmission can be measured using single and paired pulse protocols as evoked cortical responses change in presence of GABA_A_agonist^17,19,32–34^

In this study, we investigate GABAergic modulation of cortical excitability using single and paired pulses, in a circuit and cell-type specific manner using optogenetic stimulation of the pyramidal cells medial entorhinal cortex (MEC) projecting to the hippocampus (PC_MEC-> Hpc_). In addition, we characterize cortical resilience to sustained perturbations, using trains of optogenetic stimulations. In a translational approach, we show the feasibility of similar stimulation protocols to assess GABAergic transmission in one human subject with epilepsy implanted with stereo-EEG electrodes.

## Methods

### Animals

Four C57BL/6JRj male mice aged between 8 and 12 weeks old were used. Before surgeries, mice were housed in groups in individually ventilated cages, with food and water *ad libitum* under controlled conditions (12:12h light-dark cycle, constant temperature 22°C and humidity 30-50%). All mice experiments were conducted in accordance with protocols approved by the veterinary office of the Canton of Bern, Switzerland (license no. BE 19/18).

### Virus targeting

Mice were anesthetized with Isoflurane (5% in ambient air for induction and 1.5-2% for maintenance, Abbvie, Switzerland). They were then placed in a digital stereotaxic frame (David Kopf Instrument, USA) and body temperature was kept at 37°C using a rectal probe and closed-loop heating system (Harvard Apparatus, USA). Eyes were protected with ointment (Bepanthen, Bayer, Germany) and analgesia was given as subcutaneous injection of Meloxicam 2mg/kg (Boehringer Ingelheim, Switzerland). Scalp fur was removed using a depilatory cream (Weleda, Switzerland) and the scalp was disinfected with Betadine (Mundipharma, Switzerland). After skin incision, conjunctive tissues were scratched, and bur holes were drilled at the targeted locations.

To express Channelrhodopsin (Ch2R) specifically in pyramidal cells from the medial entorhinal cortex (MEC) that project to the hippocampus (PC_MEC->Hpc_), we used an intersectional strategy with two recombinant adeno-associated viruses (AAV) injected in two different target brain regions, such that only neurons transfected with both viruses would express ChR2: 1) 450nl of a mostly retrograde virus containing the opsin on inverted cassette (AAVretro_EIFa_DIO_Ch2R(H134R)_eYFP from UNC Vector Core, USA) was injected into CA1 right (coordinates: antero-posterior (AP) -2.0mm from Bregma, medio-lateral +1.3mm from Bregma and dorso-ventral -1.6mm from the skull level). 2) 450nl of a mostly anterograde virus containing the Cre recombinase under CamKII promoter to target pyramidal cells (AAV1_CamKII_Cre_SV40, Addgene, USA) was injected into the right MEC (+3.2mm laterally from Lambda along the lambdoid suture and DV -2.5mm from skull level). Viruses were loaded on a 500nl Hamilton syringe (Model 7000.5, Hamilton Company, USA) and injected using a micro-infusion pump (Pump 11 Elite Nanomite, Harvard Apparatus, USA) at the rate of 50nl/min, with 10 min pause before syringe retraction. The skin incision was then sutured and animals were put back in their home-cage. They were monitored and received analgesia (Meloxicam 2mg/kg) for 3 days.

### Mice electrodes implantation

Three weeks after viral injection, mice were implanted with skull electroencephalography (EEG) screws and intraparenchymal depth electrodes for Local Field Potentials (LFP) recordings. To obtain faster recovery after long surgeries, a reversible mix (10 μl/g) was used for anesthesia, with the following composition: 10% Midazolam 5mg/ml (Sintetica, Switzerland), 2% Medetomidine 1mg/ml (Graeub AG, Switzerland), 10% Fentanyl 0.05mg/ml (Sintetica, Switzerland) and 78% NaCl 0.9%.

Otherwise, surgery was carried out as described above through the same incision.

Bilateral frontal EEG screws (∅1.9mm, Paul Korth GmbH, Switzerland) were soldered to a stainless-steel cable (W3 wire, USA) and inserted at coordinates -1.0AP, ±2.0ML. Reference and ground EEG screws were inserted above the cerebellum and the olfactory bulb, respectively. Intraparenchymal electrodes, made of tungsten wires (∅76.2μm, model 796000, A-M System, USA), were pinned in an 18-EIB board (Neuralynx, USA), inserted one by one and glued in place at the following coordinates: MEC on the lambdoid suture, ±3.2ML, - 2.5DV; CA1 -2.0AP, ±1.3ML, -1.6DV; CA3 -2.0AP, ±2.2ML, -2.2DV and DG -2.0AP, ±1.3ML,-2.3DV. The right entorhinal electrode was glued to a homemade optical fiber implant (∅200μm, 0.39 NA Core Multimode Optical Fiber, FT200EMT inserted and glued into CFLC128 ceramic ferrules, Thorlabs, USA).

Mice were woken up with a mix for reversing anesthesia (10 μl/g), composed of 5% Atipamezole 5mg/ml (Graeub AG, Switzerland), 2% Naloxone 4mg/ml (OrPha Swiss GmbH, Switzerland), 50% Flumazenil 0.1mg/ml (Anexate, Roche, Switzerland) and 43% NaCl. After the surgery, mice were monitored in their home-cage for a week and received analgesia (Meloxicam 2mg/kg) for three days. During a brief isoflurane anesthesia, the EIB board was then connected to a HS-16-CNR-MDR50 Neuralynx cable, and the optic fiber to a home-made optical patch cord (optic fiber FT200EMT glue in a FC/PC connector, 30230G3, Thorlabs, USA). Before recordings and stimulation started, mice habituated for a week to freely move with the cables fixed to a moving hook under the same environmental conditions described above.

### Data acquisition and stimulation protocol

EEG and LFP signals were amplified and digitized at 2KHz using the Digital Lynx SX data acquisition system (Neuralynx, USA). For opto-stimulation, a patch-cord was connected to a 473nm blue laser (Cobolt 06-MLD, HÜBNER Photonics GmbH, Germany) controlled by a Matlab (Mathworks, USA) script through a pulse train generator (PulsePal 2, Sanworks, USA). The digital trigger signal was recorded along with the electrophysiology data. The analogue modulation mode of the lasers was used to stimulate with different light intensities by employing varying input voltages. Maximum intensity was calculated to be around 60mW at the tip of the optic fiber. The reliability of the laser outputs and modulation was ensured previously by recording laser power for each of the stimulation protocols with a photodiode (PM100A, Thorlabs) connected to the Digital Lynx with a Universal Signal Mouse board (Neuralynx Inc., USA).

Each of the four recording weeks included three sessions with three different pharmacological condition (Diazepam (DZ), Pentylenetetrazole (PTZ) and NaCl) interspersed with >48h in random order to allow for drug elimination and avoid excessive kindling (i.e. the tendency for seizures to become more severe over time). Each session took place in the afternoon (second half of the light phase) and lasted about 60min during which mice were kept awake by gentle handling. Diazepam (Valium 10mg/2ml i.m./i.v., Roche, Switzerland) and Pentylenetetrazole (PTZ, P6500, Sigma-Aldrich, USA) were respectively diluted at 2.5mg/ml and 10mg/ml in NaCl 0.9% in order to inject a constant volume (2μl/g i.p.) across conditions.

Each session contained 1) 10min of baseline recording, 2) a first i.p. injection (Dz, PTZ or NaCl), followed 3min later by 3) a paired pulse (PP) protocol lasting 45 min, 4) a second i.p. injection (NaCl, PTZ or NaCl, re-injecting PTZ due to its rapid elimination^23,35^, followed 3min later by 4) a single pulse (SP) protocol lasting 10min followed by 5) a seizure induction protocol.

All pulse simulations consisted of single or pairs of 3ms light pulse of different intensities delivered in a random order every 8-12 seconds with random jitter. The PP protocol lasting 45min included pairs of pulses spaced apart by 30 different inter-pulse intervals (IPI) logarithmically distributed from 6 to 2000ms. The first pulse of the pair (‘conditioning’ pulse) could vary between three different intensities (1/3 max intensity, 2/3 max intensity or max intensity), but the second pulse of the pair (‘probing’ pulse) was always set at 2/3 of the maximum laser intensity. Each of the possible IPI was presented 3 times for each of the 3 conditional pulse intensities for a total of 270 paired pulses, presented in a randomized order. The SP protocol, lasting 10min, consisted of 3ms single light pulses of 12 different intensities, linearly distributed in the range of the laser analogue modulation (0.45V to 1V) and repeated five times in random order. The first intensity corresponds to undetectable laser output and the last one to the maximum laser output intensity.

The seizure induction protocol consisted of repeated trains of 3ms light pulse at 20Hz of systematically increasing duration (1, 2, 3, 4, 5, 6, 7, 8, 10, 12, 15, 20, 25 and 30 seconds) presented at one-minute intervals. When a seizure was elicited, the protocol was stopped and the duration of the last train which elicited a seizure determined the “time to seizure” for the session. Seizures were visually detected by a trained experimenter as sustained ictal activity continuing after the end of the stimulation.

All animals (n=4) showed seizures from the first session and underwent 12 seizure inductions in 12 sessions of opto-stimulation, with 4 sessions in each pharmacological condition. One animal was excluded from LFP analysis due to lost signal in the right hippocampus.

### Mice histology

Animals were euthanized after the twelfth stimulation session. They were anesthetized with 250mg/kg Pentobarbital (Esconarkon 1:20, Streuli Pharma AG, Switzerland) and transcardially perfused first with cold NaCl 0.9% and then with 4% formaldehyde for 5min each. Extracted brains were post-fixed in 4% formaldehyde (Grogg Chemie, Switzerland) for 24h, then transferred in sucrose for 48h before being flash-freezed with -80° methylbutane. Brains were then sliced along the sagittal axes (40µm slices) on a cryostat (Hyrax C 25, Zeiss, Germany), and collected in PBS. For immunostaining against the GFP and NeuN proteins, slices were first incubated 1h in a blocking solution composed of PBST and 4% bovine serum albumin. Free-floating slices were then incubated 48h at 4° with a mix of anti-GFP (1:5000, Ref. A10262, Invitrogen, USA) and anti-NeuN (1:1000, Ref. 2931160, Millipore, USA). They were then rinsed 3×10min with PBS containing 0.1% Triton and incubated 1h at room temperature with two secondary antibodies of different colors, AlexaFluor 488 (1:500 Abacam, ab96947) for the anti-GFP primary antibody and AlexaFluor 555 (1:500, Ref. A21422, Invitrogen, USA) for the anti-NeuN primary antibody. Finally, slices were washed again 3 x 10 minutes and mounted on glass slides. Images were obtained using an epifluorescence microscope (Olympus BX 51, Olympus Corporation, JP) at different magnifications (4-10x).

### Human subject and data acquisition

Human data were collected from one medically intractable epilepsy patient undergoing invasive presurgical evaluation with stereo-EEG at Inselspital, Bern. Eight electrodes were implanted in the right hemisphere as necessary for diagnostic needs and without relationship to the present research study. These intracranial EEG electrodes enable direct cortex stimulations with pulses of electrical current to probe cortical excitability in the form of cortico-cortical evoked potentials (CCEPs). The patient provided informed consent for participation and this study was approved by the ethics committee of the Canton Bern.

### Data acquisition and stimulation protocol in humans

Each intracerebral electrode (DIXI medical, Microdeep®, France) consists of eight platinum channels with a diameter of 0.8 mm and a length of 2 mm with varying spacing. The MRI and postsurgical CT were co-registered using the Lead-DBS software (www.lead-dbs.org) to determine the exact location of each electrode contact. A neurologist (MOB) labeled the channels based on their anatomical locations. The intracranial EEG recording was amplified using a 128 channel Neuralynx ATLAS system (Neuralynx Inc., USA), with a sampling frequency of 2000Hz, a voltage range of ± 2000µV along with a digital trigger signal to identify stimulation onsets. A neurostimulator (ISIS Stimulator, Inomed Medizintechnik GmbH, Germany) was used to deliver stimulations at varying intensity.

The same stimulation protocol was repeated before and after the intravenous administration of clonazepam 0.75 mg, a GABA-A receptor agonist of the benzodiazepine class, given for medical reasons (end of clinical work-up). Although evoked responses could be identified in many electrodes, only signals from channels in the entorhinal cortex are part of this analysis, as to study a similar circuit as in mice. Individual stimulation pulses were bipolar (one pair of neighboring channels in the hippocampus) and square-biphasic (3 ms/phase). The single pulse protocol (SP) consisted of varying intensities ranging from 0.2 – 10mA, each pulse repeated three times and randomly delivered with an inter-stimulation-interval (ISI) of at least 4 s. The paired-pulse protocol (PP) consisted of a first conditioning pulse varying in intensity (1, 2 or 4mA) followed by a second probing pulse with fixed intensity of 2mA at varying inter-pulse-interval (IPI) ranging from 10 to 1600 ms (increasing logarithmically in 29 steps). The inter-stimulation-interval (ISI, between blocks of paired-pulses) randomly varied between 9-24 seconds.

### Data Pre-processing

The human LFP signals were preprocessed in Matlab (The MathWorks, Inc., Natick, Massachusetts, United States) in following steps: 1. calculating bipolar derivations by subtracting monopolar recordings from two neighboring channels on the same electrode lead, discarding most of the stimulation artifacts 2. removing 10ms of remaining stimulation artifact immediately following the trigger by linearly interpolating preceding and following mean values over 10ms, 3. bandpass 0.5 – 200Hz and 50Hz (and harmonics) notch filtering followed by resampling to a frequency of 500Hz.

The mice EEG and LFP signals were preprocessed in Python (Python Software Foundation, https://www.python.org/) with a 5-100Hz band-pass and a 50Hz (and harmonics) notch filter. For the quantification of the LFP response to light pulses, the channel from the right hippocampus with the shortest response latency upon opto-stimulation of the MEC was selected.

### Common Signal Processing and data analysis

All the signal processing and analysis were carried out using custom Python scripts.

The evoked response to a stimulation pulse was measured as the line length (LL) per millisecond (ms) of the LFP signal as follows:

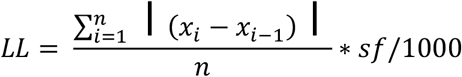

Where n is the number of datapoints over which the LL is calculated, *sf* the sampling frequency and *x* the measured local field potential at each datapoint. For single and paired pulse responses, the LL was calculated respectively over the first 100ms for mice or 250ms (precisely, 240ms of the response excluding removal of the 10ms stimulation artifact) for humans, following the onset of stimulation to account for different kinetics in mice and humans evoked responses. The window size is chosen for humans, as to include both negative peaks of a typical CCEP described in literature^36^. For the 20Hz train stimulations in mice, the LL was calculated during the 53ms window in between two pulses. For each session and each intensity, the pulse with the higher LL were visually checked to ensure that there was no artefact or removed otherwise. This process was done blind to the session condition.

After its calculation, line-length values were expressed as ratios to a reference and normalized in two different ways for each protocol. 1) For the single pulse stimulation protocol and to establish an Input/Output (I/O) curve, the LL value of each stimulation within a channel was divided by the average LL value obtained at maximal stimulation intensity during baseline recording (Respectively, 10mA stimulations in baseline condition in humans and maximum laser intensity in NaCl condition in mice). This facilitated the comparison of the varying response strength to the different stimulation intensities across sessions and subjects. 2) For the paired pulse stimulations, the LL values of both pulses are normalized by the mean LL responses to single pulse stimulation at intermediate intensity during baseline recording (2mA in humans, ⅔ laser intensity in mice), which also corresponds to the parameters of the 2^nd^ ‘probing’ pulse. Responses to the second (probing) pulse are qualified as cortical suppression or facilitation if they are greater (normalized LL > 1) or smaller (normalized LL < 1) than the reference single pulse average response. To maximize the number of trials in each analysis, the first pulse of PP stimulations with IPIs greater than the window used to calculate the LL were also included as single pulse responses.

### Statistics

Statistical testing was performed using bootstrap estimation statistics methods and graphical representations^37^. Differences between conditions are calculated and shown as the mean difference (i.e. the effect-size) and its 95% confidence interval (95%CI), obtained by performing bootstrap resampling 5000 times. To compare time to seizure across conditions, we used a two-sided sign test.

## Results

### Optogenetic probing of cortical excitability

To probe cortical excitability in a circuit and cell-type specific manner, we target pyramidal cells in the entorhinal cortex projecting to the hippocampus (PC_MEC->Hpp_) with an intersectional viral approach (Fig. 1A) which results in robust expression of Channelrhodopsin in the layer III of the medial entorhinal cortex (MEC) and in its projections to the ipsilateral Hippocampus (Hpp) (Fig1D).

**Figure 1.**
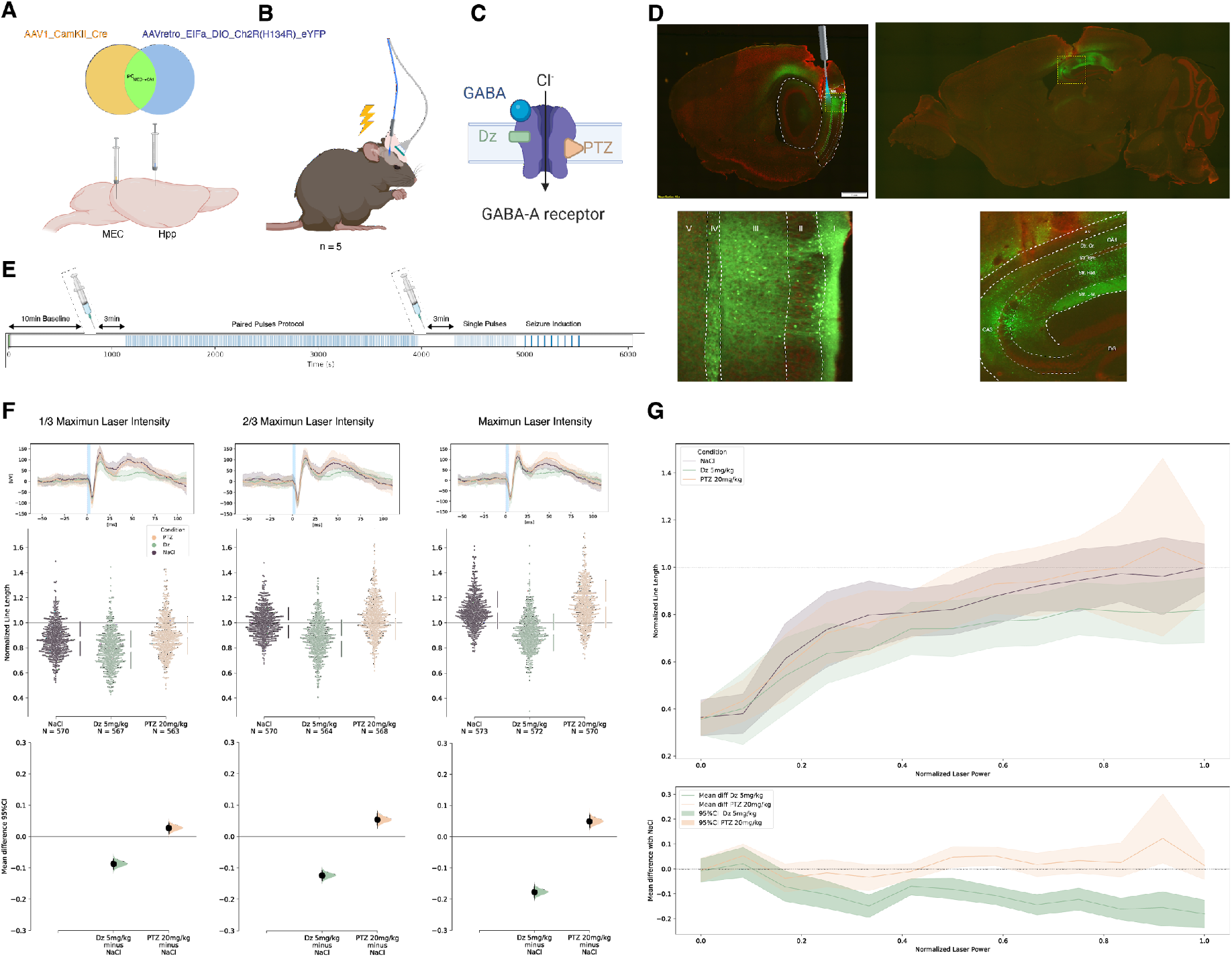
Probing limbic cortical excitability using single-pulse opto-stimulation. (**A**) Schematic of the intersectional viral approach targeting entorhinal pyramidal cells projecting to the ipsilateral hippocampus (PC_MEC->Hpp_). A retrograde AAV carrying the inverted genes for *Channelrhodopsin* (*Ch2R*) and *eYFP* in-between LoxP sites (DIO) is injected in the CA1 region of the hippocampus. A second virus carrying the gene for the *CRE recombinase* under the *CamK2* promoter (mainly expressed in pyramidal cells), is injected in the medial entorhinal cortex (MEC). Only neurons transfected by both viruses will express the opsin Ch2R and the reporter protein eYFP. (**B**) Four freely moving awake mice were stimulated with opto-pulses and recorded over 12 sessions each lasting ∼2 hours. (**C**) Diazepam 5mg/kg (Dz) and Pentylenetetrazole 20mg/kg (PTZ), which are allosteric modulators of the GABA_A_ receptor, leading to increased and decreased GABAergic transmission, respectively. A-C images created with BioRender.com. (**D**) Sagittal brain slices at 10x and close-up views (inset at the bottom) of entorhinal cortex (approximately +3.2mm laterally from Bregma) and ipsilateral hippocampus (approximately +1.3mm laterally from Bregma) revealing viral expression in the PC_MEC->Hpp_ neurons in green and ani-NeuN labeling in red. (**E**) Sequence of opto-stimulations for each session consisting of 10min baseline recording, first i.p. injection (NaCl, PTZ 20mg/kg or Dz 5mg/kg), paired pulses (PP), second i.p. injection (PTZ 20mg/kg or NaCl), single pulses (SP), and finally 20Hz pulse-trains to determine the time needed to induce a seizure (time-to-seizure, 1-30s). (**F**) Hippocampus responses to single pulses of different intensities per pharmacological conditions (color-coded) across 12 sessions shown as average LFP trace (top, mean ± SD, blue shade is 3ms light pulse), line-length (100ms LL) of individual trials (middle, N, number of pulses) and estimated mean differences (bottom, black dot) with 95% CI (bottom, black vertical bar obtained by bootstrapping). To group animals together in a swarm plot, all responses are expressed as a ratio, normalized by the individual mean response to the ⅔ maximum intensity stimulation in the NaCl session of the same week. (**G**) Average input-output response curves to SP of 12 different intensities (Top, 100ms LL mean ± SD, normalized to response to max laser intensity in NaCl condition, n∼60 pulses per condition) and estimated mean difference across pharmacological conditions (bottom, color lines and shading are 95%CI, bootstrapped).

Mice (n=4) were implanted with depth wire-electrodes in the hippocampus for LFP recording and an optic fiber in the MEC for opto-stimulation during awake and freely moving conditions (Fig 1B). Over four weeks and a total of 12 sessions, they repeatedly received a stimulation protocol to probe cortical excitability and cortical resilience.

Cortical excitability was assessed on different dimensions by measuring response to single pulse, by characterizing the input-output curve for varying stimulation intensities and by studying non-linearities in cortical response in the form of paired pulse induced cortical suppression and facilitation. Additionally, with the goal of probing cortical resilience, mice received increasingly long trains of stimulation to induce seizures and quantify the magnitude (here the duration) of the provoking perturbation necessary to reach the seizure threshold. Each week, the same protocol was done in three different pharmacological conditions: NaCl, Diazepam (Dz, 5 mg/kg) or a sub-convulsive dose of Pentylenetetrazole (PTZ, 20mg/kg).

### GABAergic modulation of evoked response to single pulse

Opto-stimulation of PC_MEC->Hpp_ with 3ms pulses induced evoked potentials in the local field potential (LFP) of the ipsilateral hippocampus after a few milliseconds as expected for direct projection (Fig 1F). In presence of diazepam, the evoked response was significantly diminished with a reduced amplitude of both the positive peak and its after-going wave (Fig 1F). PTZ showed an inverse effect with an increase in mean SP response. For both drugs, the effect was stronger for greater intensities. To quantify the difference between conditions, we calculated the line length (see methods) of the evoked response at the individual trial, on a 100ms window which included all the components of the evoked response. To characterize cortical excitability over a range of inputs, we measured an input-output (I/O) curve which was characterized by a floor effect, a relatively linear increase in response and a ceiling effect (Fig 1G). In the presence of diazepam, the excitability threshold remained unchanged, but slope of the I/O curve was decreased, resulting in increasing difference with the control NaCl condition as input intensities increased. Additionally, even at maximum stimulation intensities, responses were weaker in the diazepam conditions as compared to the NaCl condition, suggesting a subtractive decrease in response. PTZ tended to increase the slope of the I/O curve at the higher intensities (Fig. 1G, n= 60 pulses per intensity) reaching significance, albeit with small effect-sizes, when statistics were computed on sufficient amounts of data (Fig. 1F, n=570 pulses per intensity).

### GABAergic modulation of cortical facilitation

To characterize expected non-linearities in cortical dynamics, we measured cortical suppression and facilitation, using random sequences of paired pulses with 30 different inter-pulse intervals (IPI) in triplicates. The first ‘conditioning’ pulse induces a small perturbation, the lasting effects of which are probed with a second ‘probing’ pulse. For IPI between 6 and 17ms, the response to the probing pulse was decreased compared to a SP, indicating cortical suppression. Conversely, for IPI between 17 and 102ms, the response to the probing pulse was increased, indicating cortical facilitation (Fig 2A). For longer IPI (102-1995ms), response to probing pulse went back to the baseline level of single pulse response. Three different conditioning pulses were used (⅓ max laser intensity, ⅔ max laser intensity or max laser intensity) but the effects on the probing pulse were comparable and they were grouped for analysis (Fig. S2). Probing pulse was always set at ⅔ max laser intensity.

**Figure 2.**
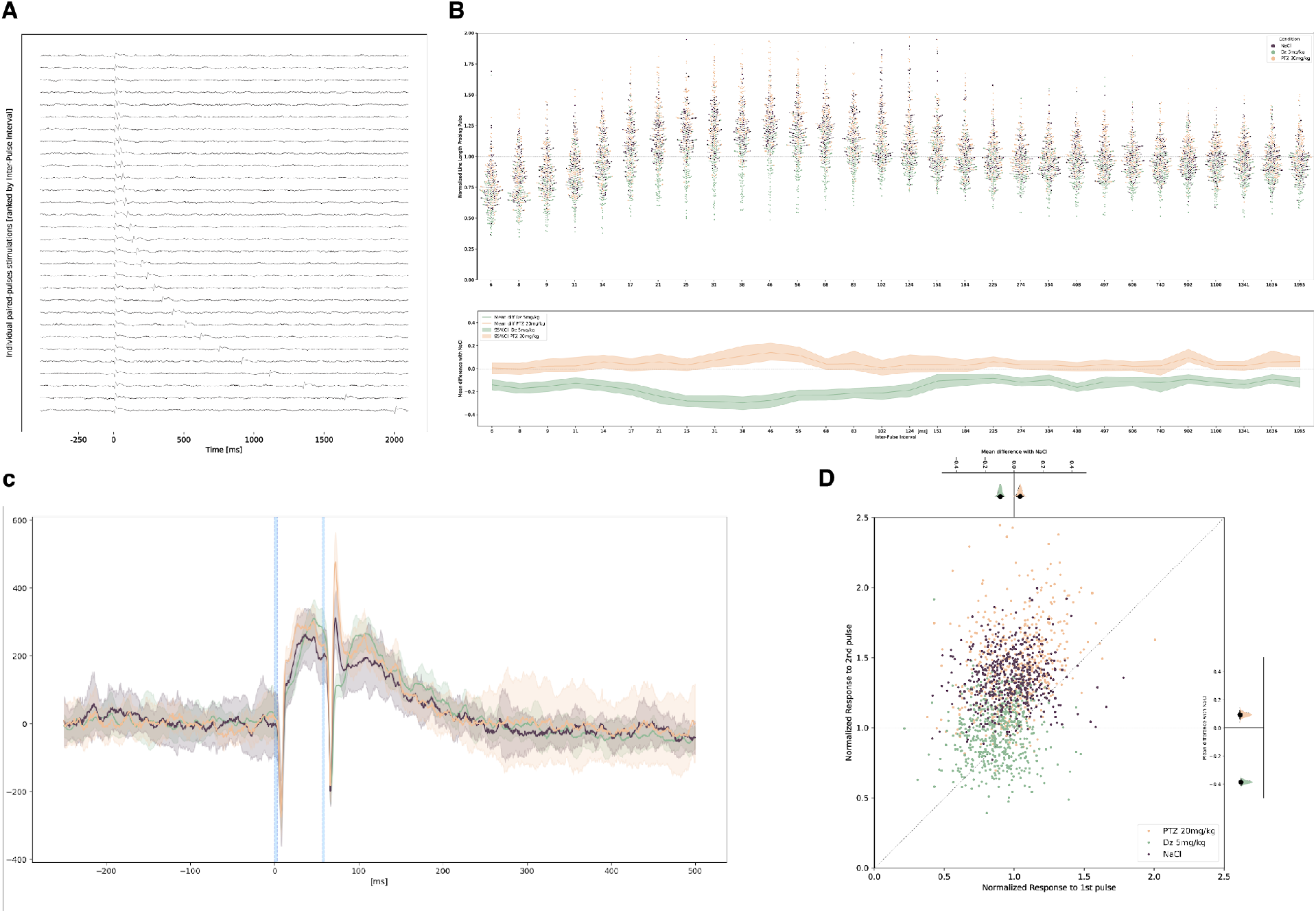
Probing limbic cortical suppression and facilitation with paired pulses. (**A**) Ranked hippocampal average LFP responses (n = 135 pulses) recorded to PP opto-stimulation of PC_MEC->Hpp_ of increasing inter-pulse intervals (IPI) in one NaCl session. Note how the second ‘probing’ pulse is reduced for IPI below 15ms (cortical suppression) but increased for IPI between 17 and 102ms (cortical facilitation) as compared to the first ‘conditioning’ pulse. (**B**) Top: Individual responses to probing pulse by pharmacological condition (color-coded), quantified as the 100ms LL, normalized to the mean response to SP in the NaCl condition and plotted as a function of the IPI with the preceding conditioning pulse. *Bottom panel* is showing the corresponding estimated mean difference with the NaCl condition and its 95%CI (shading, bootstrapped). Cortical facilitation is visible for IPI between 27 and 102ms for the NaCl and PTZ conditions but is greatly decreased in presence of Dz. (**C**) Average LFP trace (±SD) for paired-pulses with a 56ms IPI for each condition, in one representative animal (4 sessions, 36 3ms-pulses for each condition, blue shading). (**D**) Post-hoc analysis of paired pulses in the facilitation range (31 to 68ms). For individual paired pulses, the response for both pulses is plotted as the LL on a 31ms window, normalized to the mean SP response in the NaCl condition. Cortical facilitation is visible for the NaCl condition (points above the diagonal), increased for PTZ and lost for Dz (in average centered on diagonal), resulting in greater mean differences for response to second pulses compared to the first ones (top and right panel, black line and shading are 95%CI and distribution, bootstrapped).

In the presence of diazepam, cortical suppression was increased, and cortical facilitation was strongly decreased (Fig 2B). The mean difference between NaCl and Dz was maximum for facilitation IPI range (17-102ms), indicating a specific effect on cortical facilitation. For IPI longer than 102ms, differences between NaCl and Dz was similar to the one observed in response to SP (Fig 1F). On the LFP trace, this decreased response to PP was visible as a complete reduction of the positive peak (Fig. 2C).

To confirm this specific effect on facilitation, we performed an additional post-hoc analysis by evaluating the responses to both pulses from pairs with IPI leading to the largest facilitations (31 to 68ms). For NaCl, the responses to second pulses were typically greater than to the first ones whereas for Dz both pulses had similar responses (centered on the diagonal, Fig. 2D). This was confirmed by greater mean differences between second pulses than between first pulses. PTZ showed an opposite modulatory effect, with an increased facilitation (Fig 2B-D) but a lack of effect on suppression. These differences were also visible as a higher positive peak in the LFP trace (Fig 2C). Evoked cortical facilitation and suppression were stable across sessions and weeks (Fig S3).

### Optogenetic probing of cortical resilience

In a dynamical system, an increase in excitability should correspond to a more unstable state and therefore a decreased distance to a critical transition^4,5,12,38^. To test this assumption, we quantified the amount of perturbation necessary to provoke a seizure, using 20Hz trains of opto-stimulation with increasing duration (1s to 30s) until the induction of a seizure. The time of opto-stimulation necessary to induce a seizure -”time to seizure”-was used as an indicator of cortical resilience. As previously shown for principal cells in the CA1 subfield of the hippocampus^23,39^, optogenetic sustained stimulation of PC_MEC->Hpp_ was sufficient to elicit limbic seizure in non-epileptic mice (n= 4 out of 4). During the train stimulations, LFP response in the right hippocampus showed spike-shaped discharges time-locked to the opto-pulses (Fig 3A and B). This entrainment was immediately visible on the ipsilateral (right) hippocampus but began later on the contralateral hippocampus (Fig. 3D). Ictal changes were seen at the end of the seizure-inducing train of pulses and consisted of rhythmic high amplitude spikes (at lower frequency than stimulation), sometimes followed by low voltage fast activity (Fig. 3A, B and D). Opto-induced self-sustained seizures typically lasted for 20s after the end of the train stimulation.

**Figure 3.**
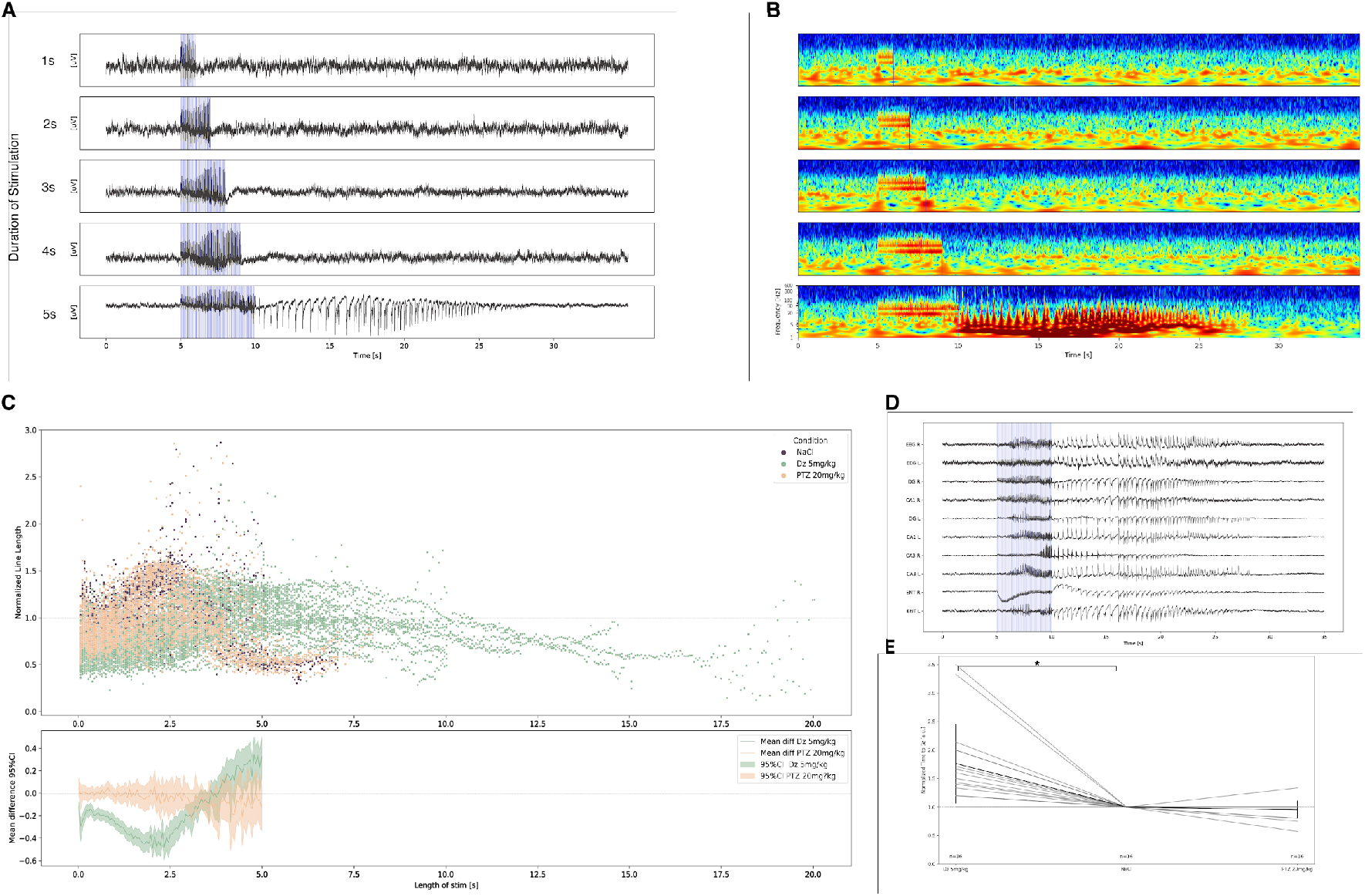
Probing limbic cortical resilience with train opto-stimulations. (**A**) Series of right hippocampus LFP traces in one animal undergoing the seizure induction protocol with trains of 20Hz opto-stimulations (blue vertical lines) of increasing duration until a seizure is induced, here after 5s stimulation in a NaCl session. Examples for Dz and PTZ are shown in Fig S3. (**B**) Wavelet spectrogram of the LFP traces in *A*. For the trains of stimulation which did not provoke a seizure, only the frequency band corresponding to the stimulation and its harmonics are visible (here, 20 and 40Hz). Black vertical lines indicate the beginning and end of the train stimulations. During the last stimulation train, which provokes the seizures, ictal activity is visible as a mix of lower (∼10Hz) and higher-frequencies (>100Hz) in the spectrogram, and is thereafter self-sustained. (**C**) *Top:* individual responses to each 3ms light pulse (color-coded per pharmacological condition), quantified in the right hippocampus LFP as 53ms LL, normalized to the mean response in the NaCl condition. The response increases gradually during the train until 2.5s. This increase is slower in the Dz condition, reflecting decreased facilitation. *Bottom:* estimated mean difference with the NaCl condition and its 95%CI (shading, bootstrapped) calculated when 20 trains of both conditions were available (<5s). (**D**) Recording of the optogenetically induced seizure in *A* (5s) across the 10 recording channels (EEG R: frontal right EEG screw, EEG L: frontal left EEG screw, DG: dentate Gyrus, CA1: Cornu Ammonis Area 1, CA3: Cornu Ammonis Area 3, ENT: medial entorhinal cortex). (**E**) GABAergic modulation of “time to seizure”, measured as the time of 20Hz optogenetic stimulation of the PC_MEC->Hpp_ necessary to induce a seizure and normalized (within each stimulation week) to the NaCl condition. Time to seizure increases in presence of Dz (p < 0.001, Sign test). No difference was found for the PTZ condition.

### GABAergic modulation of cortical resilience

Next, we examined whether GABAergic modulation of cortical excitability could correlate with changes in cortical resilience, measured as the time-to-seizure. The 20Hz frequency was chosen because it corresponds to the IPI showing cortical facilitation with PP, and where the effect of GABAergic modulation was stronger (50ms, see Fig 2B). During train stimulations, we observed a build-up of the facilitation during the first few seconds (0-2.5s), where the responses to opto-pulses became increasingly strong (Fig 3C). In the presence of Dz, the cumulative increase in pulse responses was slower between 0 and 2.5s, consistent with the loss of facilitation observed in Fig 2B-D. For PTZ, no difference in response to train stimulation was observed compared to NaCl.

These observations correlate with longer trains of stimulations necessary to induce a seizure in the Dz condition. When normalized to the NaCl session of the corresponding week, Dz session needed between 1x and 3.5x longer stimulations to induce a seizure (mean ± SD: 1.76 ± 0.70, p < 0.001 Sign-Test). No significant difference in time to seizure was found for PTZ (mean ± SD: 0.95 ± 0.15, p = 0.076 Sign-Test).

### Modulation of cortical excitability in one patient with epilepsy

To confirm the ability of single and paired pulse to probe cortical excitability in humans, we used a series of bipolar electrical stimulation of the hippocampus and measured generated responses in four electrodes situated in the ipsilateral entorhinal cortex of one patient who had previously been implanted with stereo-EEG for a pre-surgical investigation of their intractable epilepsy (Fig 4A). During the last day of clinical investigation, the patient received an intravenous administration of clonazepam 0.75 mg, a GABA-A receptor agonist of the Benzodiazepine (BDZ) class with similar effects as diazepam. Before and after the injection, protocols of SP and PP analogous to the ones used in mice were carried out.

**Figure 4.**
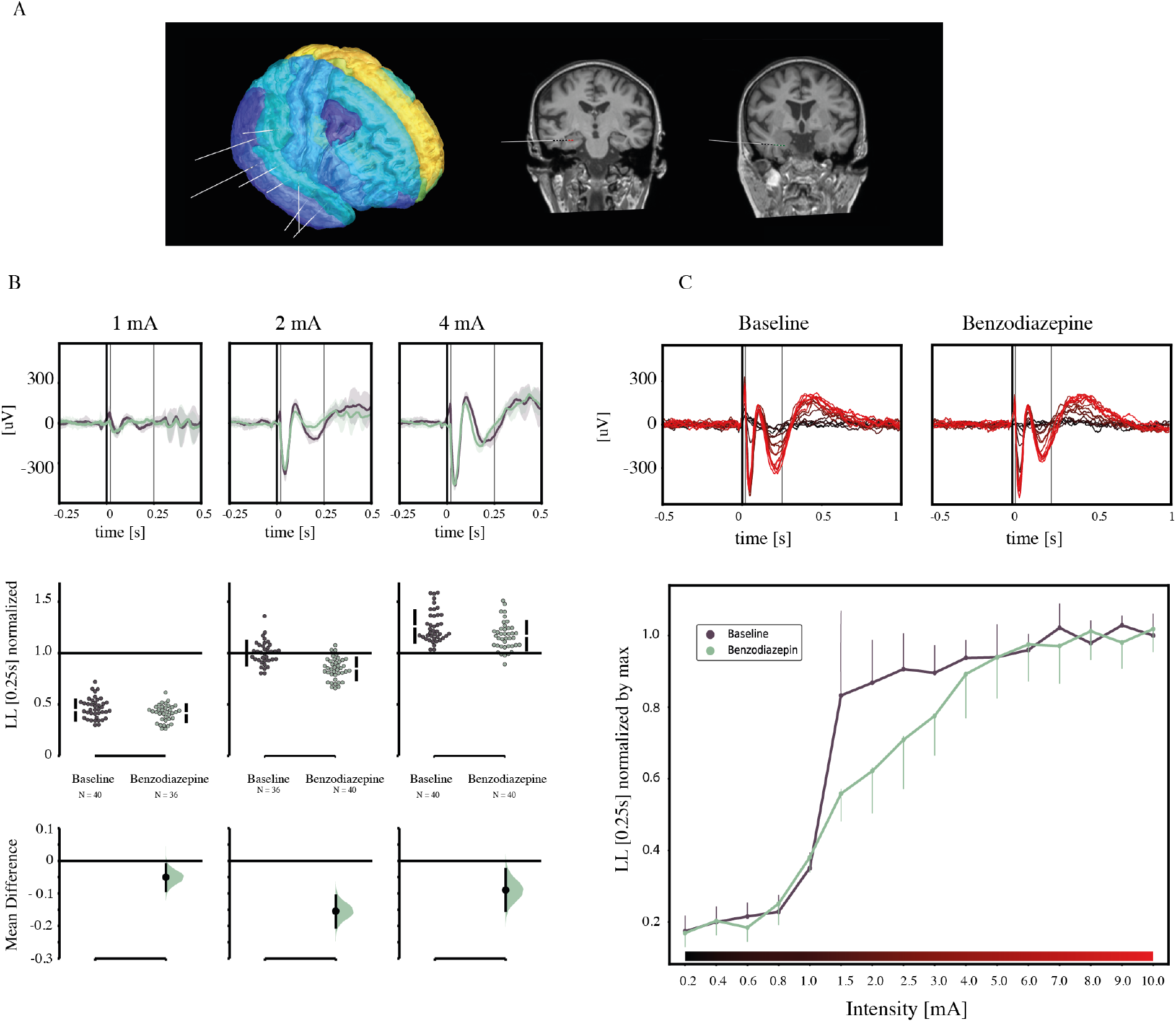
Probing limbic cortical excitability with single electrical pulses in one patient with epilepsy. (**A**) 3D-reconstruction of full implant (left) and coronal MRI slices with the location of the stimulation channels in the hippocampus (red, middle) and the recording channels in the entorhinal cortex (green, right). (**B**) Top: Average LFP (shading is ±SD) in one response channel for each conditioning intensity (left to right) and for baseline (brown) and after benzodiazepine injection (green). Thin black vertical lines are the borders of the window for LL calculation [10ms, 250ms]. Thick black vertical line is the onset of the stimulation. Middle: Individual responses in 4 channels, along with average (±SD, black vertical bars) quantified as 240ms LL and normalized within channels by the average 2mA baseline response. Bottom: estimated mean difference (black dot, bootstrapped effect size) and 95% CI (black vertical lines) of the benzodiazepine effect for three stimulation intensities. (**C**) Measure of cortical excitability as a function of the input intensity. Top: Average LFP responses in one channel to increasing stimulation intensities (color-coded, black: 0.2mA, bright red: 10mA) for both conditions (left: baseline, right: benzodiazepine). Bottom: quantification of the average LL across four channels (dots,±SD) normalized within channels by the average response to a 10mA stimulation.

Single square-shaped biphasic electrical stimulations (3ms per phase) produced evoked responses in the local field potential that lasted longer (∼700-800ms) than the opto-pulse response in mice (∼100ms) (Fig 4B). In the presence of clonazepam, the response to single pulse was significantly reduced for intensities from 1-5mA, with a maximum difference at 2mA and a decrease in slope (Fig 4B-C, 95% CI via estimation statistics). As for opto-stimulation evoked responses in mice, the predominant BDZ effect seems to be on the second, slower, part of the evoked response (compare Fig. 1F and 4B). Unlike in the mice experiment, the maximum evoked response could be attained at stronger stimulations, despite the presence of clonazepam. The discrepancy concerning the reaching of the I/O curve plateau between optogenetic and intracranial electrical stimulation could be explained by different factors such as cell-type specificity of the optogenetic stimulations or differences in cortex organization across species and brain structure. However, one simple explanation may lie in the fact that the range of explored input intensities for the optogenetic stimulation wasn’t high enough to get to this plateau. In support of this hypothesis, SP intensities with the maximum mean difference between benzodiazepine and the control conditions correspond to max intensity for optogenetics simulations but only half maximum intensity of the electrical ones (Fig 1F and 4B).

To parallel characterization in mice of non-linear cortical responses, the PP protocol was composed of pairs with 30 different IPIs and three different intensities of conditioning pulses (1, 2 or 4mA). All probing pulses were set at 2mA.

For conditioning pulses of 1mA, the IPIs between 6 and 38ms induced cortical suppression, whereas IPIs between 38ms and 500ms led to cortical facilitation (Fig 5A). Compared to mice findings, the range of IPI for cortical suppression and facilitation were both shifted toward longer intervals (compare Fig. 2B and 5A). This difference could be due to differences in stimulation methods, species or length of stimulation (3ms and 6ms respectively) and evoked response.

**Figure 5.**
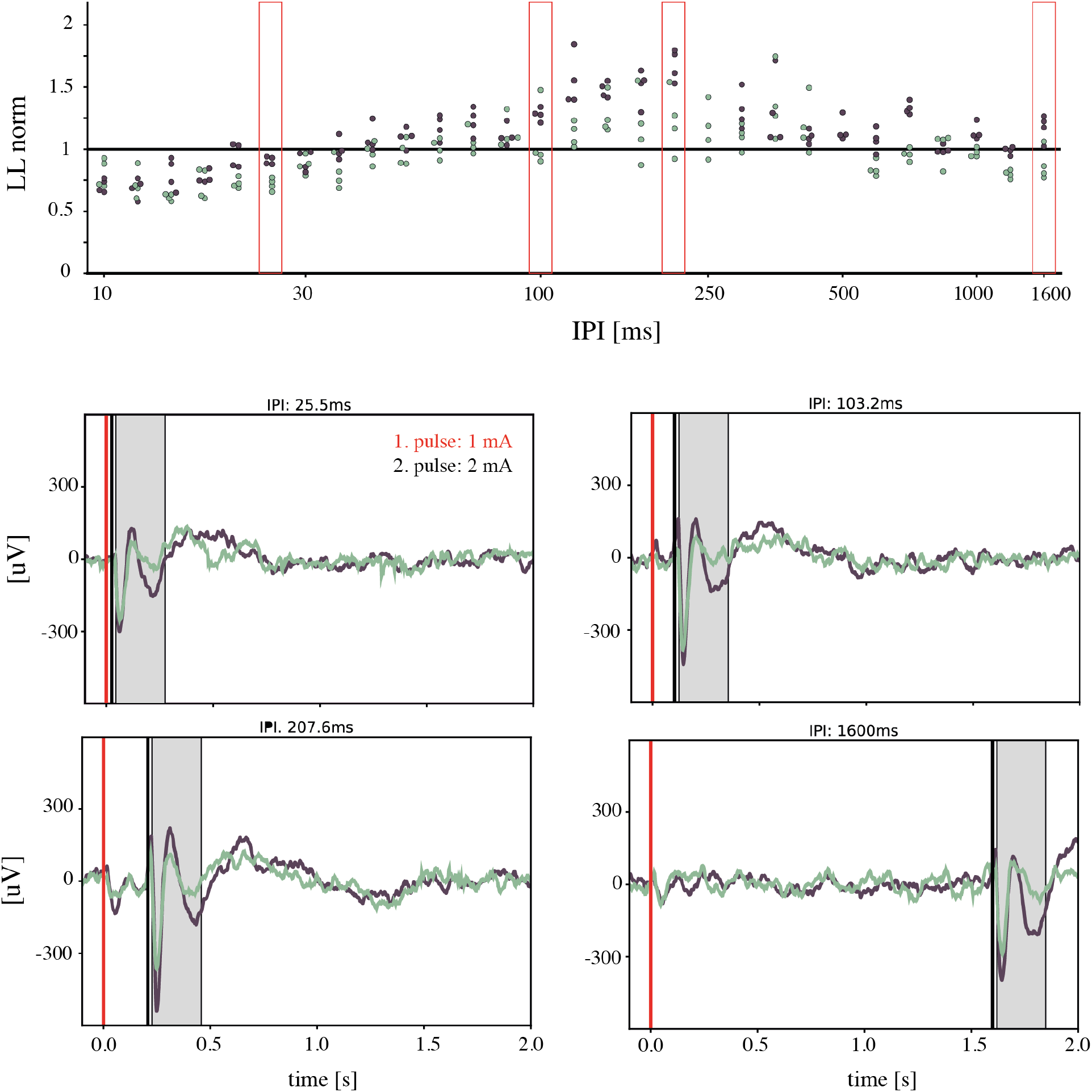
Probing limbic cortical suppression and facilitation with paired-pulses in one patient with epilepsy. Individual entorhinal cortex responses to a probing pulse (2mA) after 1mA conditioning pulse (2mA and 4mA conditioning pulses in supplementary Fig. 5) by pharmacological condition (color-coded, black is baseline, green after benzodiazepine injection) in four channels, quantified as the 250ms LL and normalized within channels by the average response to a 2mA SP in the baseline condition. Red boxes are shown at the bottom as the LFP for single trials examples in one channel for four PP stimulations of different inter-pulse-intervals (IPI: 25.5, 103.2, 207.6, 1600ms). Grey block indicates the window for calculating the LL.

In the presence of clonazepam, cortical facilitation was reduced as previously observed for opto-stimulations in mice (Fig 2B). For IPIs corresponding to cortical suppression, the BZD effect seems comparable to differences in SP. Due to low numbers of data points per conditions, no estimation statistics were done. For other conditioning pulses intensities (2 and 4mA), this effect was lost and no difference between BZD and baseline was observed (Fig S5).

## Discussion

Here, we characterized a range of cortical dynamics across two species and found a correlation between different degrees of cortical excitability and the resilience to ictal transitions. In mice, we used a circuit and cell-specific approach to drive excitatory pyramidal neurons of the limbic cortex, increasing the specificity of our findings. In a translational approach in humans, we used electrical pulses to probe different aspects of cortical excitability in homotopic circuits, supporting the translational potential of our findings. Specifically, we showed that GABAergic modulation of cortical resilience was detectable by changes in responses to single and paired-pulses, providing a proof-of-principle that changes in cortical excitability can be tracked with simple probing protocols. The changes in single and paired pulse response observed in our study have never been, to our knowledge, directly correlated with actual changes in seizure susceptibility.

From a dynamical system perspective, repetitive stimulation can be seen as a perturbation which forces the system to pass the seizure threshold, an unstable tipping point, beyond which self-sustained seizures are unavoidable^4,5,11,12,38^. In-silico simulations predict that increasing levels of cortical excitability decrease the distance to the seizure threshold and therefore the amount of perturbation needed to tip the system into a seizure state^4,11^. These concepts are in line with previous work mostly carried out *in-vitro*^4,11^. Validation of these relationships *in-vivo* requires tools to probe both cortical excitability and resilience. To that aim, we developed a circuit-specific seizure model together with protocols to probe cortical excitability and measure the ‘time-to-seizure’, the duration of a set perturbation necessary to induce a seizure, keeping stimulation frequency, pulse width and intensity constant. Changes in single and paired pulse evoked responses observed in presence of benzodiazepine correlated with actual changes in ‘time-to-seizure’, confirming the link between cortical excitability and resilience in freely moving mice. Going beyond the unnatural conditions of seizing brain slices maintained in low ion concentrations, our methods provide means to measure cortical excitability and resilience in a freely moving non-epileptic animal, opening the way to a fundamental understanding of the mechanisms governing the fluctuations of cortical excitability *in-vivo*. Indeed, our model has the key advantage of not requiring any kindling and induces seizures on the first attempt, in an otherwise intact network.

Crucially, in addition to measuring resilience to seizure induction, we also developed two protocols of minute perturbations to probe cortical excitability without inducing irreversible state transitions. Specifically, in the presence of a GABA_A_-R agonist in both animals and one human, we found a decrease in the slope of input-output curves, and an even stronger decrease in cortical facilitation, along with increased resilience measured only in mice. These effects were somewhat variables on a single trial basis, but stable on average within and across stimulation sessions. Cortical facilitation for inputs at 30-100ms interval may play an important role in reducing resilience, as both optogenetic and electrical induction of seizure in CA1 seem to be most effective between 10 and 30Hz^22,40,41^. In fact, cumulative facilitation was visible in the trains of pulses preceding the induction of seizures, resulting in the build-up of increasingly stronger evoked responses over a few seconds. In the presence of benzodiazepine, the build-up rate decreased corresponding to longer time-to-seizure. This observation suggests a common mechanism for cortical facilitation and loss of resilience, although further studies are needed to confirm this. Any synchronous entrainment of neuronal firing may result in a seizure, including trains of optogenetic stimulations in the hippocampus^22,23^ and in the neocortex of rodents^24,39^, as well as electrical or magnetic repetitive cortical stimulation on non-epileptic human brains^8,9^. At the cellular level, several explanations have been proposed such as accumulation of extracellular potassium^40,42^, switch to excitatory GABA after intracellular chloride accumulation^43^ or depolarization block of the inhibitory interneurons.

Probing cortical excitability with short perturbations has been applied in a number of research settings, and may emerge as a clinical modality. Evoked cortical responses to single or paired stimulation had been previously measured in the presence of GABAergic drugs by TMS-EEG^19,33,44,45^. These studies systematically found that diazepam and other benzodiazepines decreased cortical facilitation^32^ in line with our findings. Although these methods are helpful to carry out noninvasive measurements in human subjects, they lack cell type and circuit specificity. In presurgical epileptology, where intracerebral electrodes are implanted for clinical reasons, the level of circuit specificity can be refined, and has been advantageously used for the localization of epileptic parenchyma^13^. To our knowledge, the opportunity to characterize drug effects on cortical excitability in this patient population has not been exploited yet, and our inclusion of one human patient in this study thus represents a first. In animals, the recent development of optogenetics adds cell-specificity to the approach of probing cortical excitability. Klorig et al.^23^ showed in a recent study, that single optogenetic pulses could be used as a measure of the cortical excitability threshold, defined as the laser light intensity at which an LFP response was measurable 50% of the time, akin to the chronaxie in older electrophysiology studies^46^. In this work, thresholds of cortical excitability were related to thresholds of seizure induction, measured as the laser intensity necessary to induce a seizure over a fixed duration of stimulation. Despite, the different metrics to characterize cortical excitability (threshold vs. input-output curves and paired-pulse facilitation) and seizure thresholds (resilience to stimulation intensity vs. time), our results are in agreement, confirming that measuring the response to small perturbations is a good proxy to evaluate cortical resilience. Our approach may have the advantage of not relying on probabilistic thresholds, which require several trials for their determination.

Cortical gain regulation, typically measured as input-out (I/O) relationships between a sensory stimulus and the firing rate of a neuron, is a fundamental property of the brain as it allows neurons to adaptively modulate their sensitivity to sensory inputs without losing their selectivity^47^. Here, we characterized I/O relationships to cortical inputs at the population -as opposed to individual neuron - level by measuring the evoked response in LFP trace produced by different stimulation intensities. In the presence of benzodiazepine, the I/O curve was shifted to the right, but the excitability threshold remained unchanged. At the neuronal level, such right shifts are typical of a decrease response gain mediated by GABAergic inhibition^48,49^. Thus, the observed right shift at the population level could correspond to lower response of individual neurons or lower numbers of recruited neurons for similar stimulation intensities.

The biological mechanisms underlying cortical suppression and facilitation are poorly understood but are believed to depend on both cell and circuit properties and are not reducible to short-term synaptic plasticity^50^. Cortical facilitation is believed to be the resultant of balance between NMDA-R dependent excitation and GABA inhibition^51,52^ and could be a sensible marker of cortical excitability. Paired pulses studies on epileptic cortices typically find increased cortical facilitation^20,53^ and have been used to track epileptogenesis^54^ but the common mechanisms linking cortical facilitation and seizure susceptibility still remains unknown.

Our study has some limitations. First, the recorded local field potentials result from the superposition of extracellular voltage potentials from different cell types and neuronal components (axons, dendrites, soma)^55^. Thus, cellular mechanisms governing changes in cortical excitability observed here cannot be inferred from our study. Still, our study bears specificity at the mesoscale, as we targeted mono-synaptic connections in both the human and mice experiment to explore cortical excitability, an approach that is more specific than many previous studies in humans. Better characterizing the link between neuronal and cortical excitability will require future work. Second, we use a model of repetitive stimulation which could lead to seizures by different ways than those taken by spontaneous seizure. Nevertheless, the high specificity with which cortical resilience can be determined is likely to be relevant to spontaneous seizures in epilepsy. Indeed, determining seizure thresholds with methods less specific than ours has been the workhorse of pharmacological developments in epilepsy for decades^56,57^. Third, cortical excitability tested under Pentylenetetrazole showed minimal, although significant changes and no change in time-to-seizure. This could relate to the very short effects (<15min) of sub-convulsive doses of Pentylenetetrazole.

In conclusion, our study provides *in-vivo* experimental evidence in support for the dynamical systems point-of-view proposed to formalize notions of cortical excitability and resilience. Taking advantage of a novel optogenetics model of ‘seizures on demand’, we propose methodology to delineate the landscape of physiological and pathological cortical excitability, including a quantification of seizure thresholds. Such tools, here piloted in one human, open the possibility of characterizing changes in cortical excitability as a function of endogenous factors or pharmacological interventions. Finally, with the fast development of closed-loop neurostimulation devices for the treatment of neurological^58–62^ or psychiatric conditions^63^, tractable markers of cortical excitability may crucially inform the choice of stimulation parameters in the future.

## Conflict of interest

none.

## Acknowledgements

none.

## Supplementary Materials

**Supplementary figure S1.**
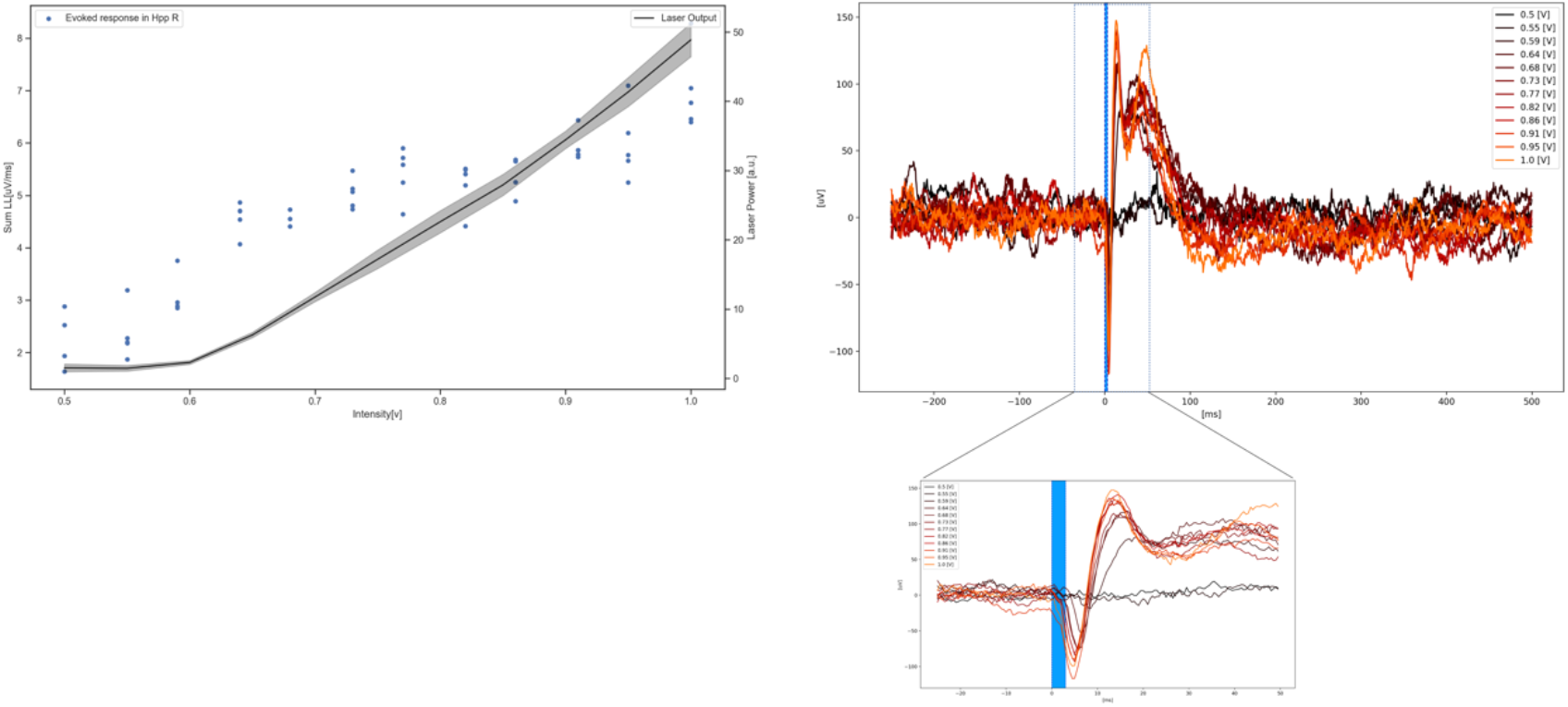
LFP response in the right hippocampus as a function of laser intensity. (A) Measured laser power (a.u. left y-axis) and LFP response (LL, right y-axis). Note the non-linear relationship between laser output and neural response, measured as the local field potential. (B) Mean LFP response for each of the 12 laser intensities used in the study. *Bottom panel* in a close-up view of the blue box.

**Supplementary figure S2.**
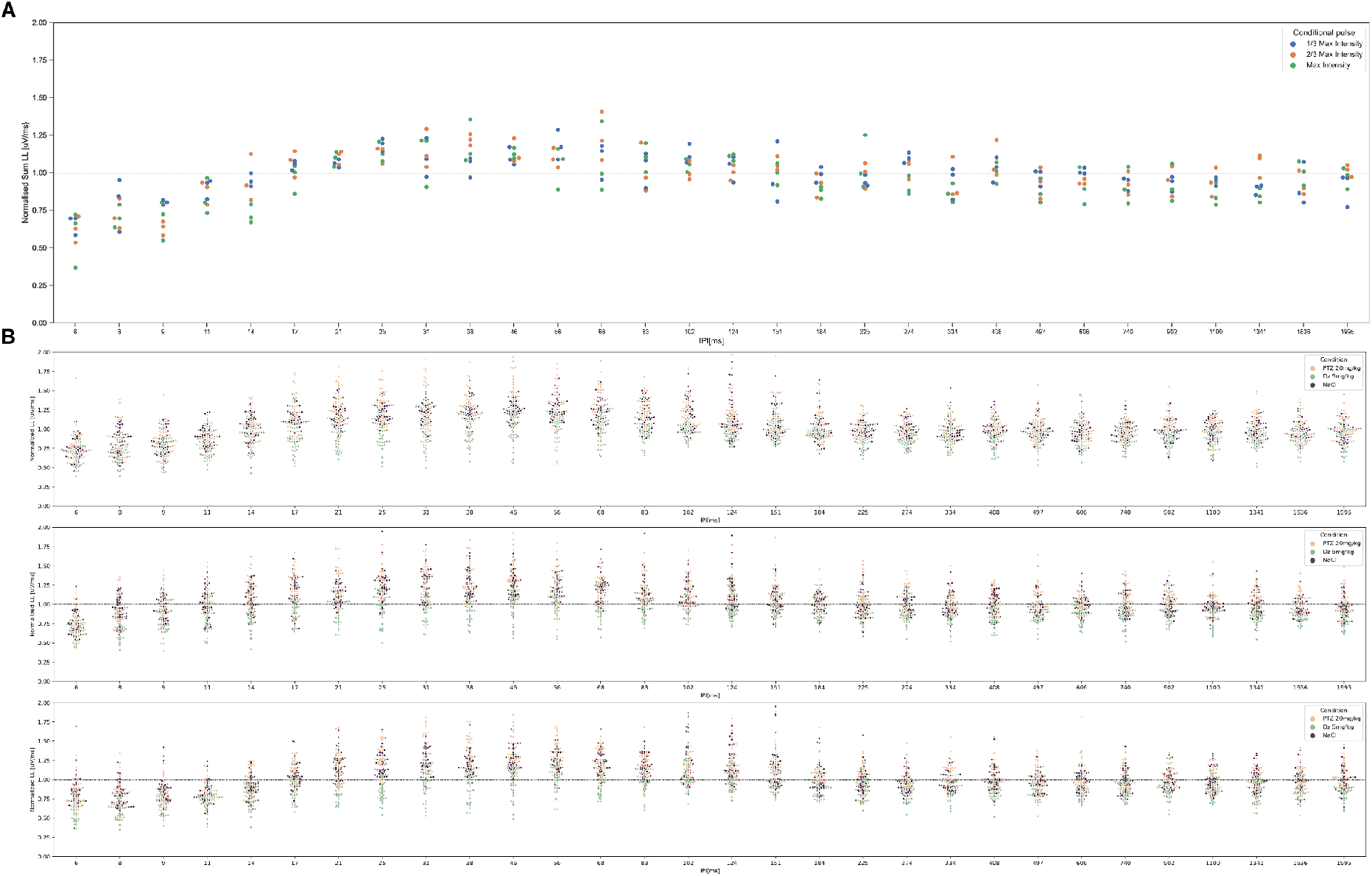
Effect of the conditional pulse intensity on the probing pulse response. (**A**) Example of probing pulse responses in a NaCl session. Probing pulses were always set at ⅔ max intensity but three different conditioning pulses were used: ⅓ max laser intensity, ⅔ max laser intensity or max laser intensity. Overall no difference due to conditioning pulse intensity was visible and they were grouped for figure 2B. (**B**) Quantification of the response to the probing pulse across pharmacological conditions for each conditioning pulse intensity. *Top panel* shows probing pulse responses after conditional pulse of ⅓ max laser intensity, *Middle panel* for conditioning pulses of ⅔ max laser intensity and *bottom panel* for conditioning pulses of max laser intensity The panel shown in Fig. 2B is the combination of all three panels.

**Supplementary figure S3.**
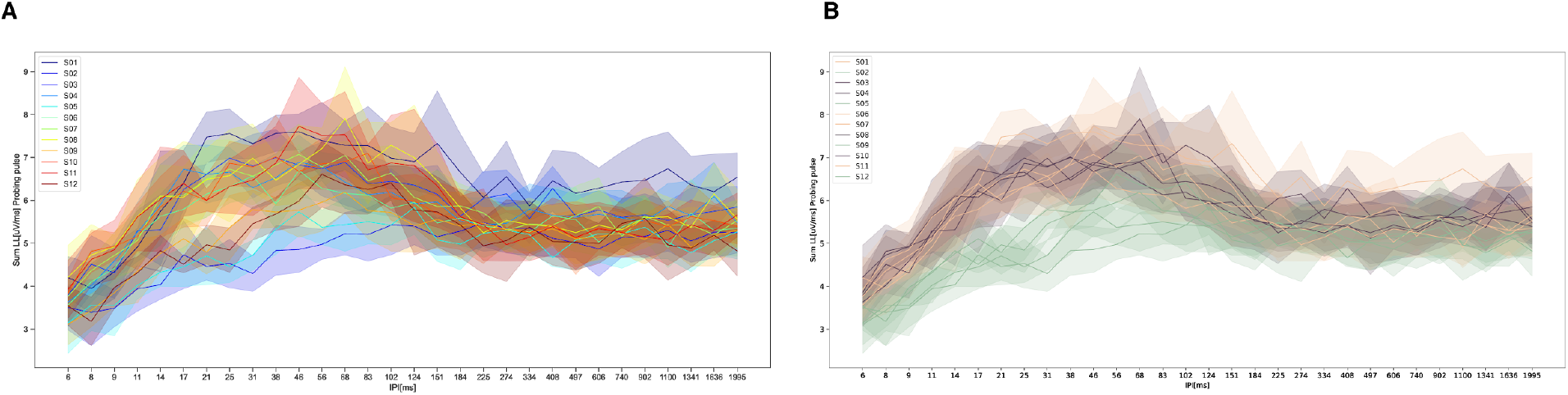
Paired-pulse cortical facilitation and suppression over all sessions. (**A**) Mean probing pulse response in function of the inter-pulse intervals, in one representative animal and for the twelve sessions. Color-code by session number, shadow areas are standard deviation. (**B**) Same graph than A but color-coded by pharmacological condition. Cortical facilitation and suppression were stable across sessions and weeks, but consistently modified by GABAergic modulation.

**Supplementary figure S4.**
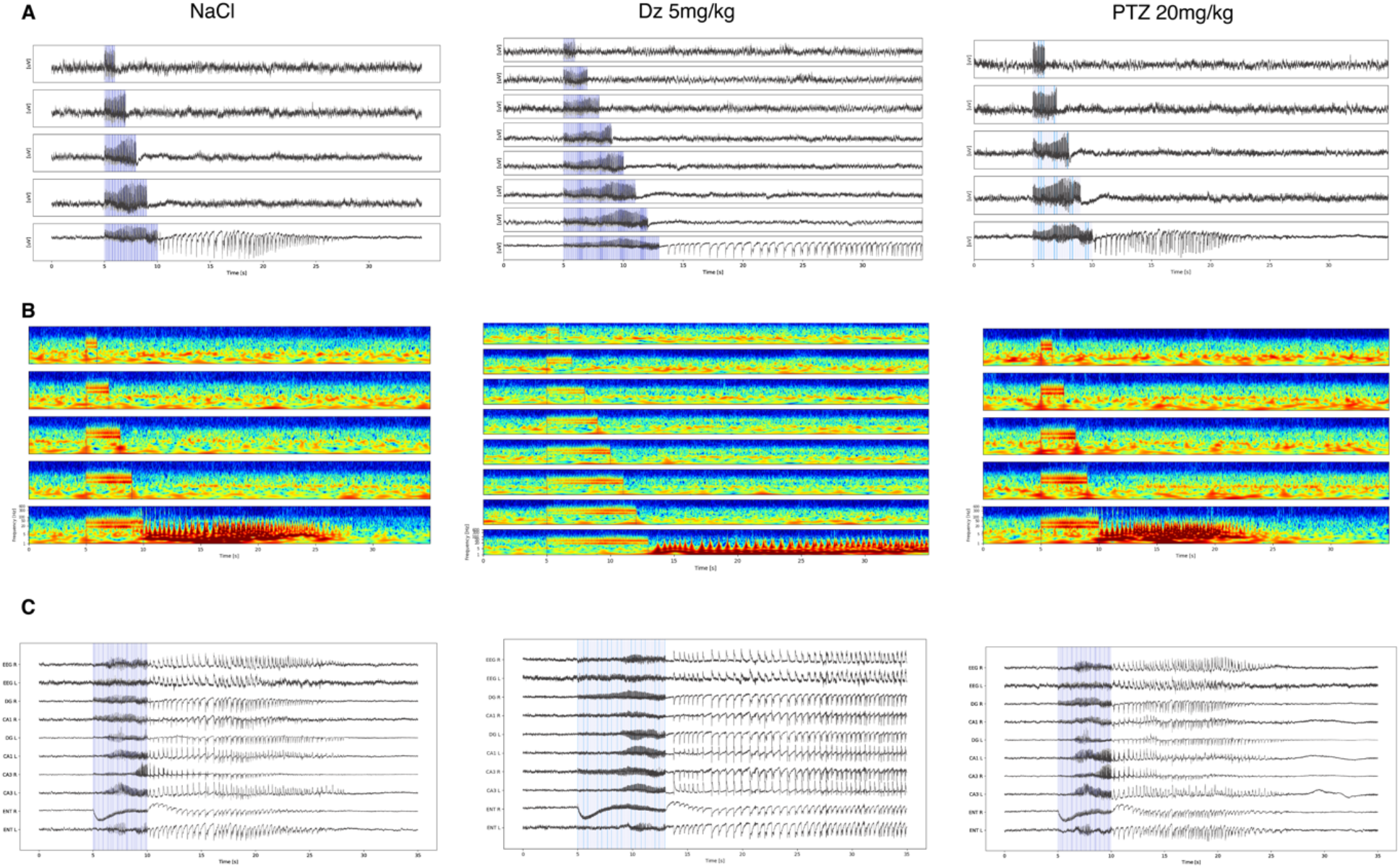
Example of seizure induction protocol for the three different conditions in the same week in one animal. (**A**) LFP trace of the right hippocampus during the 20Hz stimulation. (**B**) Time-frequency spectrograms of the trace in *A*. (**C**) Seizures recordings in 10 channels. EEG R: frontal right EEG screw, EEG L: frontal left EEG screw, DG: dentate Gyrus, CA1: Cornu Ammonis Area 1, CA3: Cornu Ammonis Area 3, ENT: medial entorhinal cortex)

**Supplementary figure S5.**
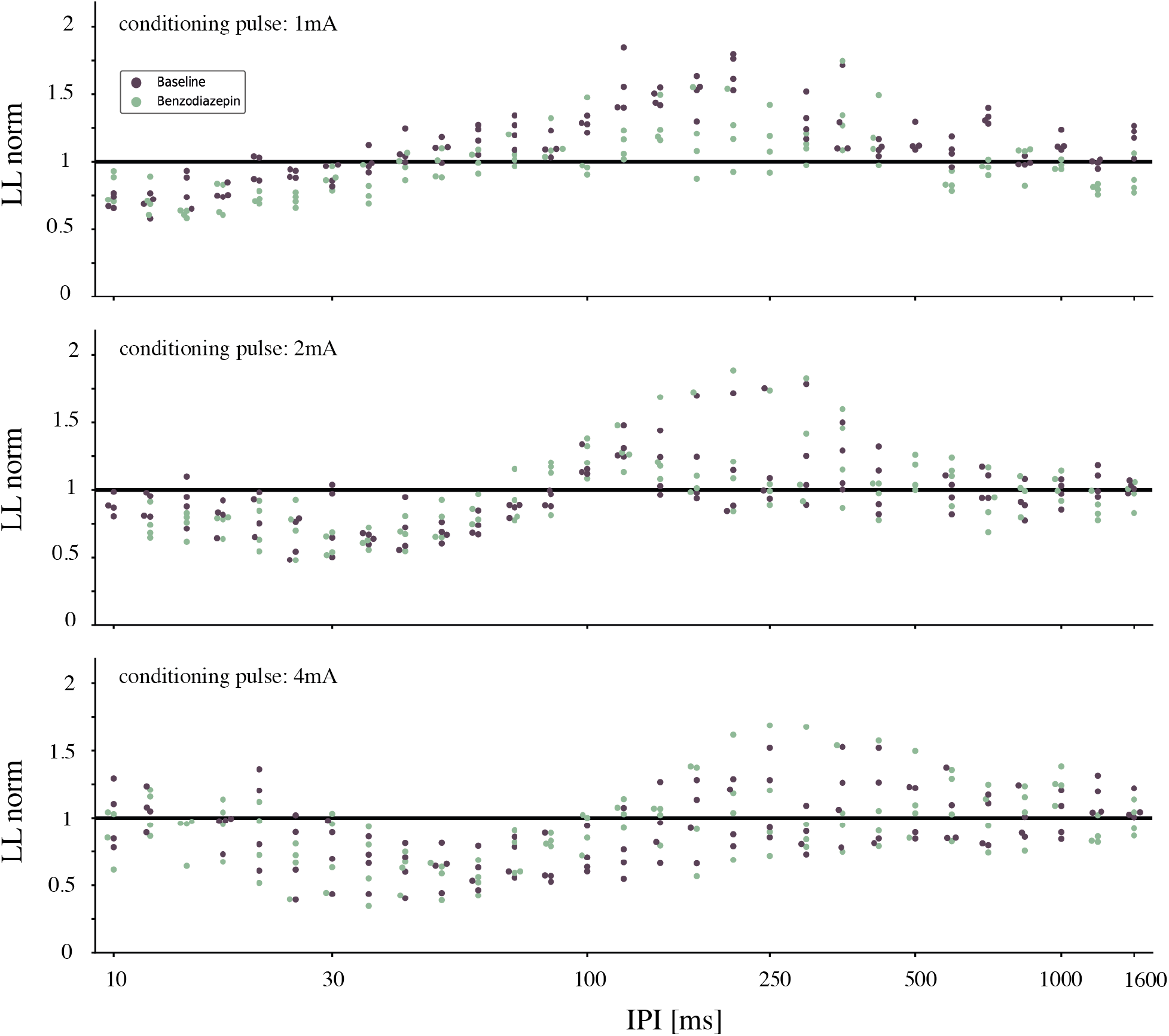
Effect of the conditional pulse intensity on the probing pulse response for intracortical electrical stimulations. LL [250 ms] to the probing pulse (2mA) after 1mA (top), 2mA (middle) and 4mA (bottom) conditioning pulse, normalized by the reference (mean LL of a 2mA SP baseline stimulation) for each IPI across the four response channels in the Entorhinal cortex. The condition is color-coded (purple: during baseline, green: during benzodiazepine).

## Notes

### Competing Interest Statement

The authors have declared no competing interest.

